# A Simple Strategy for Identifying Conserved Features across Non-independent Omics Studies

**DOI:** 10.1101/2023.11.22.568276

**Authors:** Eric Reed, Paola Sebastiani

## Abstract

False discovery is an ever-present concern in omics research, especially for burgeoning technologies with unvetted specificity of their biomolecular measurements, as such unknowns obscure the ability to characterize biologically informative features from studies performed with any single platform. Accordingly, performing replication studies of the same samples using different omics platforms is a viable strategy for identifying high-confidence molecular associations that are conserved across studies. However, an important caveat of replication studies that include the same samples is that they are inherently non-independent, leading to overestimating conservation if studies are treated otherwise. Strategies for accounting for such inter-study dependencies have been proposed for meta-analysis methods devised to increase statistical power to detect molecular associations in one or more studies. Still, they are not immediately suited for identifying conserved molecular associations across multiple studies. Here, we present a unifying strategy for performing inter-study conservation analysis as an alternative to meta-analysis strategies for aggregating summary statistical results of shared features across complementary studies while accounting for inter-study dependency. This method, which we call “adjusted maximum p-value” (AdjMaxP), is easy to implement with inter-study dependency and conservation estimated directly from the p-values from each study’s molecular feature-level association testing results. Through simulation-based assessment, we demonstrate AdjMaxP’s improved performance for accurately identifying conserved features over a related meta-analysis strategy for non-independent studies. AdjMaxP offers an easily implementable strategy for improving the precision of analyses for biomarker discovery from cross-platform omics study designs, thereby facilitating the adoption of such protocols for robust inference from emerging omics technologies.

## 1 Introduction

Omics assaying technologies are continuously advancing the types of biomolecules that can be profiled at a high-throughput scale. However, a caveat to these burgeoning technologies is that their reliability may be unvetted, and a “gold standard” has yet to be established, particularly in terms of the specificity of their molecular capture and susceptibility to technical artifacts. False discovery has been an ever-present concern in omics research. It is commonplace for potential discoveries of omics research to undergo additional validation, generally by being supported by analysis of independent omics data or by targeted in-vivo experiments. Proteomics is one such omics domain for which multiple novel platforms have been recently developed, promising improved throughput, replicability, and efficiency for measuring protein abundance compared to liquid chromatography-tandem mass spectrometry, which has been the dominant platform (Cui et al., 2022). Emerging technologies include SomaScan (Lollo et al., 2014) and Olink platforms, currently designed to measure the abundance of ∼7,000 and ∼3,000 human proteins, respectively. However, the specificity of their respective protein capture is unclear, and studies have reported a lack of concordance between these platforms (Haslam et al., 2022; Katz et al., 2022). Accordingly, these unknowns obscure the ability to characterize biologically informative features from studies performed with any single platform.

Performing replication studies of the same samples using different omics platforms is a strategy for identifying high-confidence molecular associations. This approach, which we call inter-study conservation, prioritizes associations that demonstrate statistical significance across studies. The most straightforward strategy for this approach is to perform independent analyses across studies and establish conserved associations based on shared statistical significance. However, this strategy is prone to loss of sensitivity, especially for omics studies in which multiple hypothesis correction procedures are generally warranted. In contrast, meta-analyses generally seek to increase the sensitivity of feature discovery by aggregating statistical results across complementary studies without requiring that associations reach statistical significance within each study. While this approach gains statistical power, it is susceptible to spurious findings driven by artifacts of any one study, especially without post-analytical consideration.

An important caveat of replication studies that include the same samples is that they are inherently non-independent. In this context, inter-study conservation needs to address the statistical dependency of the results from analyzing the same samples using different technologies. Otherwise, treating the studies as independent overestimates the effective number of studies performed, leading to inflation of type 1 error. Province and Borecki (Province and Borecki, 2013)proposed a strategy for accounting for inter-study dependencies based on the correlation between statistical results across studies. This strategy assumes that most statistical results arise from the null hypothesis for omics studies. Thus, the correlation between inter-study results captures the degree of their statistical dependency. However, Province and Borecki’s (Province and Borecki, 2013) implementation of this strategy is an adaptation of Stouffer’s Z-score meta-analysis method ⍰ and thus suffers from complications in establishing inter-study conservation.

Here, we present a simple statistical method for performing inter-study conservation analysis as an alternative to meta-analysis strategies for aggregating summary statistical results of shared features across complementary studies. Our strategy adapts that of Province and Borecki to account for inter-study dependency. Thus, our method functions as a complementary strategy for aggregating summary statistical results across studies that reduces the influence of artifacts in any study, thereby reducing the potential for reporting spurious findings.

## 2 Methods

### 2.1 Identifying conserved statistical associations for shared features across correlated data sets

Our approach, which we call **“adjusted maximum p-value” (AdjMaxP)**, is a strategy for aggregating inter-study statistical testing results to identify features for which evidence of statistical association is conserved while accounting for inter-study dependency arising from shared subjects. AdjMaxP is performed using vectors of nominal p-values calculated from feature-level, e.g., gene-level and protein-level, association testing results from individual studies, and aggregates multiple results for each shared feature to a single p-value. This approach offers a significant advantage over the “naïve” approach of assigning conservation based on shared statistical significance across studies. Accordingly, high-confidence features can be identified, even for instances where either study does not meet statistical significance criteria. Moreover, by accounting for inter-study dependency, our approach controls the type 1 error rate, otherwise inflated when assuming independence.

AdjMaxP builds on the maximum p-value test proposed by Wilkinson(Wilkinson, 1951). Suppose we have *N* studies, each reporting association p-values (*p*_*i*1_ … *p*_*iK*_) for *K* omics features. We assume that the results from the *N* studies are correlated, for example, if samples from the same subjects are used with different technology. Wilkinson’s maximum p-value test calculates an aggregate p-value, *P*_*agg*_, for each feature *k* directly as:

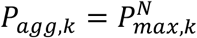

where

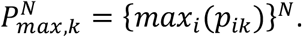

Intuitively, if the maximum p-value is small, we can reasonably infer consistent association across all studies. However, *P*_*agg,k*_ assumes independence across the *N* studies, and inflation of the type 1 error will occur when this assumption is violated. Accordingly, we devised the AdjMaxP to better control the type 1 error rate when studies are non-independent. The intuition of the method is such that if study results are correlated, then the “effective number of studies” (ENS) is smaller than *N*. In the extreme case of perfect inter-study dependence, each *N* p-values, *p*_*ik*_, must be equal, and the ENS equals 1. The opposite situation is when the studies are independent, and ENS=*N*. In general

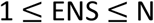

and an estimate of the aggregate p-value 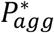 that is inter-study dependency corrected is given by

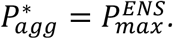

To determine an adequate estimate of ENS, we modified the Stouffer’s Z-score-based meta-analyses of large-scale omics studies introduced by Province and Borecki (Province and Borecki, 2013) that accounts for inter-study dependence. For each feature *k*, Stouffer’s Z-score aggregates the study-specific z-scores

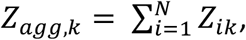

where *Z*_*agg,k*_ follows a normal distribution, *N*(0, *N*), under the null hypothesis. When the study results are correlated, Province and Borecki demonstrated that *Z*_*agg,k*_, follows a normal distribution, *N*(0, *V*) with *N* ≤ *V* < *N*^2^, and V is calculated by summing all values of the N by N matrix of pairwise inter-study correlation,

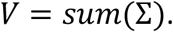

For “omics” studies, they propose estimating Σ directly from the *N* vectors of study-specific z-scores that are calculated as the compliment probit transformation of the p-values, and Σ is estimated using tetrachoric correlation after transforming z-score values into two categories defined by z > 0 and z < 0. As justification for tetrachoric correlation over Pearson correlation, they argue that for feature-by-feature hypothesis testing of omics data sets, most statistical tests follow the null distribution, such that features under the alternative hypothesis inflate estimation of the inter-study dependency if measured by Pearson correlation, leading to reduced statistical power. Under this assumption, tetrachoric correlation is preferable because it dampens the leverage of these features.

To estimate ENS, we implement a method, first proposed by Li et al. (Li et al., 2012, 2011), which utilizes the eigenvalues, *λ*_*i*_, following eigen-decomposition of Σ, and is calculated by

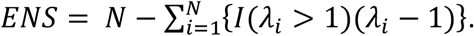

Under inter-study independence, all eigenvalues are equal to 1, yielding an ENS = N. Under complete inter-study dependence, *λ*_1_ = *N*, yielding an ENS = 1, satisfying 1 ≥ ENS ≥ N.

### 2.2 Simulation-based evaluation of validation method

To assess AdjMaxP’s performance in identifying conserved statistical associations under inter-study dependence, we performed a simulation study by repeatedly simulating N correlated sets of *K* z-scores. In these simulations, the z-scores represent compliment probit transformation of p-values from study-specific significance tests. Simulations and analyses were performed using the R statistical programming language (v 4.1.0).

For each simulation, we generated subsets of the *K* features under the null hypothesis of no association *S*_*n*_, an unconserved subset *S*_*u*_ of features that were significant in only one study, and a conserved subset *S*_*c*_ of shared true positives across the N studies. Each simulation was initialized by simulating P = 10,000 sets of z-scores from a multivariate normal distribution with mean = 0 and covariance given by an N-dimensional correlation matrix, where off-diagonal correlation values represent the background level of inter-study dependence. Unconserved and conserved subsets of features, *S*_*u*_ and *S*_*c*_, were simulated by adding a signal value, S, to initialized z-scores. For *S*_*u*_ features, signal was added to between 1 and N-1 studies; for *S*_*c*_ features, signal was added to all N studies. To evaluate the performance under different conditions, the simulated data sets were generated under various combinations of parameters. These parameters include number of studies, N; the inter-study background correlation level, C; the level of signal, S, added to either *S*_*u*_ and *S*_*c*_ features; and the proportion of the total features, Pr, represented by either *S*_*u*_ and Sc. The specific values of these evaluated included

- N: 2, 3
- C: 0, 0.1, 0.2, 0.3, 0.4, 0.5, 0.6, 0.7, 0.8, 0.9, 1.0
- S: 0, 2, 4
- Pr: 0.0, 0.2.

Simulations based on each combination of parameters were performed 25 times. Multivariate normal distributions were simulated using the *mvrnorm()* function in the MASS R package (v7.3-58.1).

For comparison, we evaluated both AdjMaxP and the Province and Borecki method. We applied these methods for each simulation and calculated a series of performance metrics based on the discovery of Sc features at both nominal p-value and Benjamini-Hochberg false discovery rate (FDR) corrected q-value (Benjamini and Hochberg, 1995) thresholding. Finally, model performance was evaluated based on specificity, sensitivity, precision, and area-under-the-curve (AUC) measures.

## 3 Results

To evaluate the relative performance of AdjMaxP and the Province-Borecki method for identifying features for which statistical associations are conserved across studies under inter-study dependency, we ran these methods on simulated test results representing varying levels of inter-study dependency, and unconserved and conserved statistical association. The distributions of performance estimations demonstrated little variability across replicate simulations of the same conditions. Mean performance estimates across all metrics and conditions evaluated are reported in Supplementary Table S1 and Supplementary Table S2 for two- and three-study simulations, respectively.

We first assessed the specificity of these methods in simulations in which no features are conserved based on a nominal p-value threshold of 0.05. Under this condition, if no unconserved features were present in the data, both methods performed similarly and consistently yielded specificity estimates near the expected of 0.95 across all background correlation levels (Figure 1 Ai, Bi). However, for simulations with 20% unconserved features, the performance of the two methods diverged, with AdjMaxP consistently yielding higher specificity across all background correlation levels (Figure Aii-iii, Figure Bii-iii). Moreover, trends of AdjMaxP were consistent across different magnitudes of unconserved signal, while Province-Borecki yielded markedly decreased specificity between signal magnitudes of 2 (Figure Aii, Bii) and 4 (Figure Aiii, Biii).

**Figure 1:**
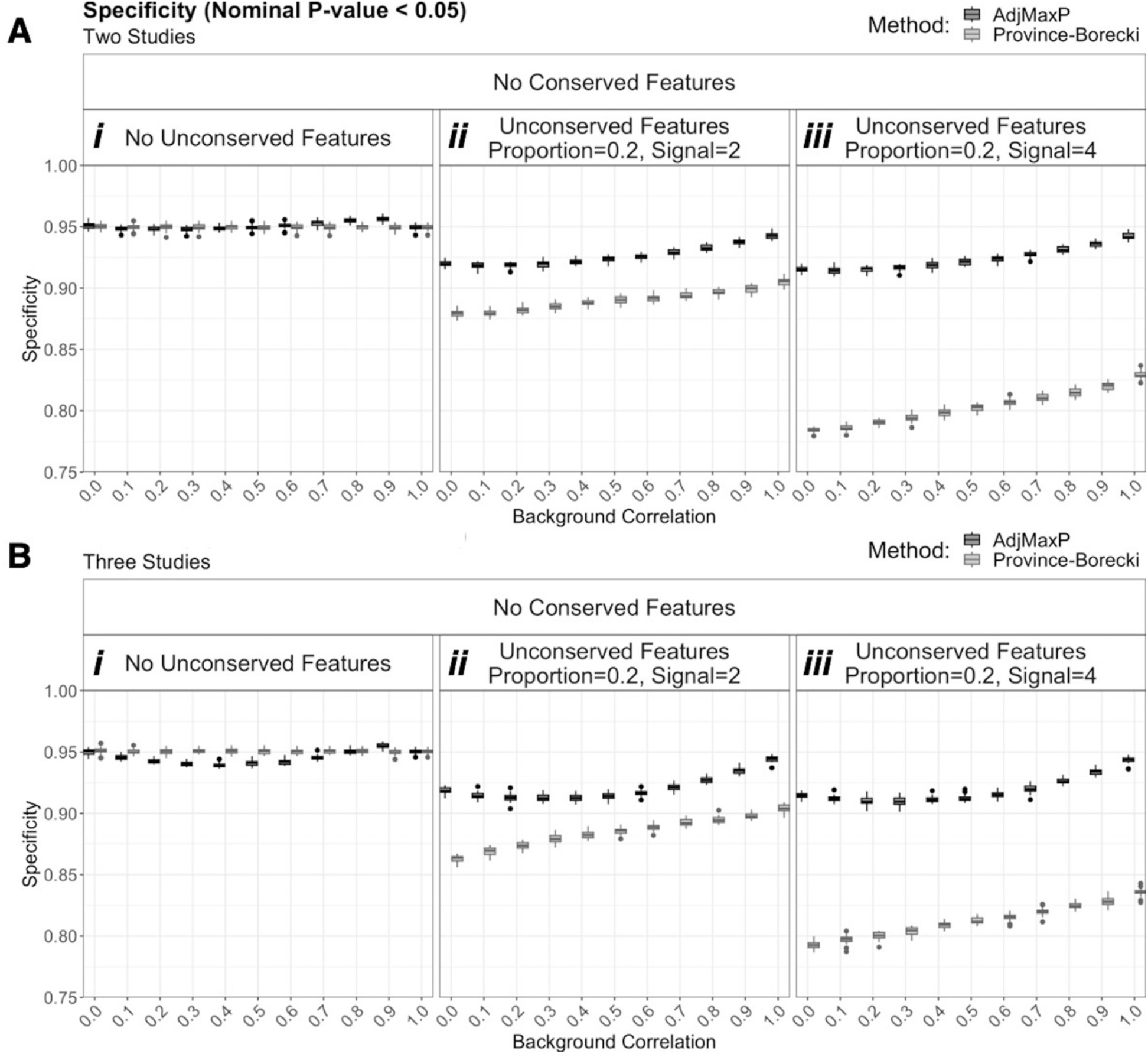
Comparison of specificity simulations of two and three studies with no conserved features, based on a nominal p-value thresholding of 0.05. Boxplots reflect the distribution of specificity at p-value threshold, 0.05, for 25 simulations of 10,000 shared features at different levels of background correlation and unconserved signal, across simulations of two and three studies with no conserved features. “Conserved” and “Unconserved” features refer to features for which signal was added (i.e. deviate from the null distribution) across all studies or fewer than all studies, respectively. “Proportion” and “Signal” indicate the proportion of features for which signal was added and the magnitude of added signal, respectively. A) Two studies. Full simulation results are shown in Supplementary Table S1. B) Three studies. Full simulation results are shown in Supplementary Table S2.

Next, given that in-practice feature discovery is generally performed based on statistical significance thresholding, we evaluated these methods’ relative precision, specificity, and sensitivity based on FDR-corrected q-value thresholding of 0.05. At this significance threshold, the precision of these methods diverged under 20% unconserved features and as the magnitude of unconserved signal increased with AdjMaxP outperforming Province-Borecki (Figure 2A, Supplementary Figure S1A). Notably, at high unconserved signal magnitude, the precision of Province-Borecki was especially low, yielding mean estimates of ∼0.55 for two-study and ranging between 0.60 and 0.50 for three-study simulations. In contrast, all mean precision estimates of AdjMaxP were greater than 0.85. (Figure 2Aiii, Supplementary Figure S1Aiii). Finally, These trends in precision were related to those of sensitivity and specificity, and AdjMaxP generally yielded higher specificity and lower sensitivity compared to Province-Borecki (Figure 3B-C, Supplementary Figure S2B-C).

**Figure 2:**
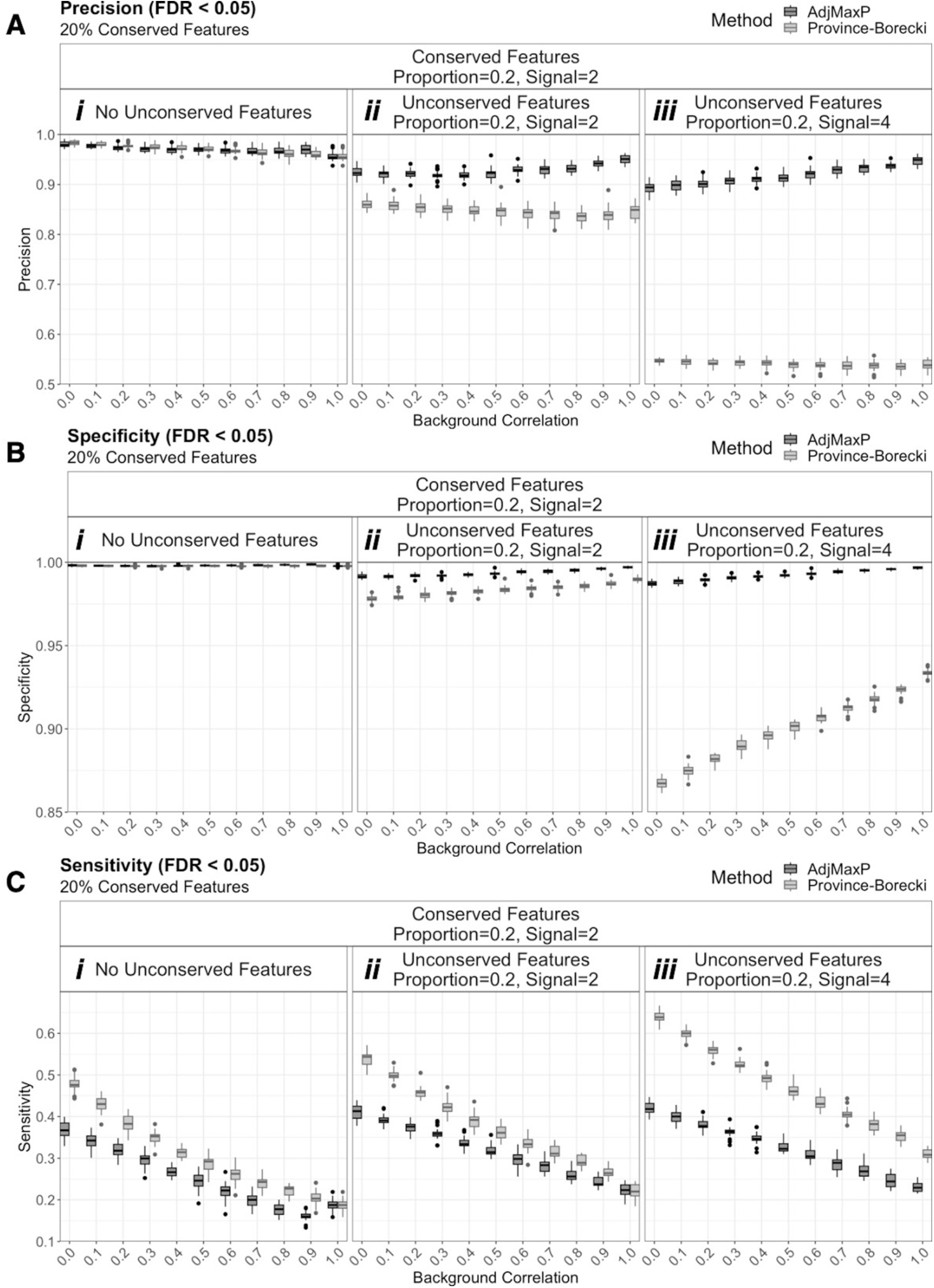
Comparison of precision, specificity, and sensitivity from simulations of two studies with 20% conserved features, based on an FDR corrected q-value threshold of 0.05. Boxplots reflect the distribution of performance at FDR q-value threshold, 0.05, for 25 simulations of 10,000 shared features at different levels of background correlation and unconserved signal, across simulations of two studies with 20% conserved features. “Conserved” and “Unconserved” features refer to features for which signal was added (i.e. deviate from the null distribution) across all studies or fewer than all studies, respectively. “Proportion” and “Signal” indicate the proportion of features for which signal was added and the magnitude of added signal, respectively. Full simulation results are shown in Supplementary Table S1. A) Precision B) Specificity C) Sensitivity

**Figure 3:**
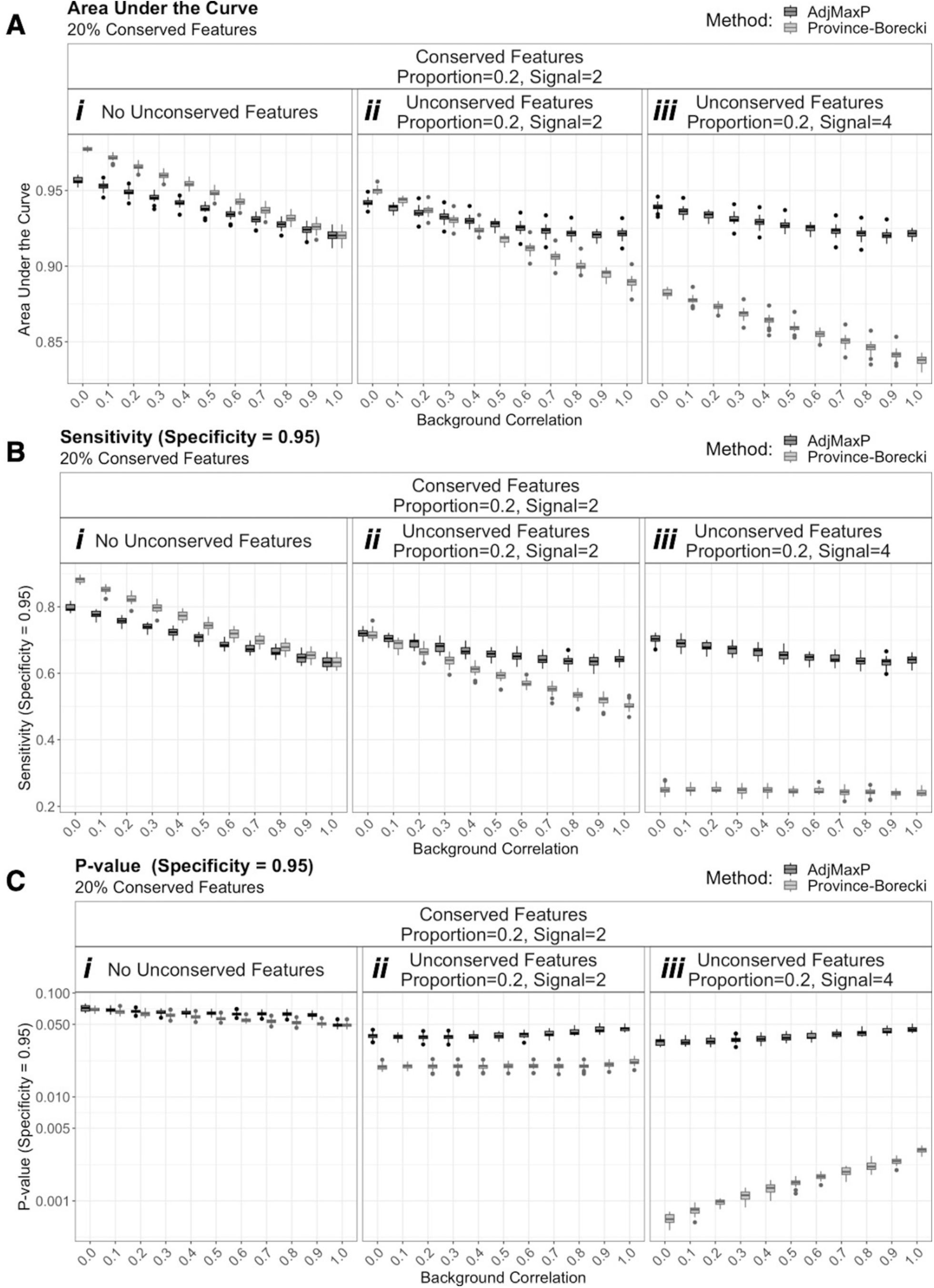
Comparison of performance under threshold variable performance from simulations of two studies with 20% conserved features, based on an FDR corrected q-value threshold of 0.05. Boxplots reflect the distribution of performance for 25 simulations of 10,000 shared features at different levels of background correlation and unconserved signal, across simulations of two studies with 20% conserved features. “Conserved” and “Unconserved” features refer to features for which signal was added (i.e. deviate from the null distribution) across all studies or fewer than all studies, respectively. “Proportion” and “Signal” indicate the proportion of features for which signal was added and the magnitude of added signal, respectively. Full simulation results are shown in Supplementary Table S1. A) Area under the curve. B) Sensitivity of conserved feature identification when specificity is equal 0.95. C) Nominal p-value threshold when specificity is equal to 0.95.

Finally, we evaluated the relative performance of these methods across varying significance thresholds using AUC, as well as sensitivity, given a set specificity threshold of 0.95. For either metric, the relative performance of AdjMaxP and Province-Borecki varied based on the presence and magnitude of unconserved signal (Figure 3, Supplement Figure 2). If no unconserved features were present in the data, Province-Borecki generally outperformed AdjMaxP (Figure 3Ai, Bi, Supplementary Figure S2Ai, Bi). However, at 20% unconserved features of high signal magnitude, 4, AdjMaxP outperformed Province-Borecki (Figure 3Aiii, Biii, Supplementary Figure S2Aiii, Biii). Finally, at lower unconserved signal magnitude, 4, the performance of the two methods was similar, and AdjMaxP outperformed Province-Borecki at higher levels of background correlation (Figure 3Aii, Bii, Supplementary Figure S2Aii, Bii). This difference in sensitivity between methods can be attributed to differences in the p-value threshold that yields 95% specificity. For AdjMaxP, at 20% unconserved features, the p-value required to achieve 95% specificity remains ∼0.05, while for Province-Borecki this p-value threshold drops to below 0.005 for high unconserved signal (Figure 3Cii-iii, Supplementary Figure S2 Cii-iii).

Overall, AdjMaxP outperforms Province-Borecki under conditions in which unconserved features are present in the data across various metrics, and the level of divergence increased with the magnitude of unconserved signal. Moreover, trends in AdjMaxP performance estimates demonstrated minimal changes across varying magnitudes of unconserved signal. Finally, these trends were consistent across simulations of two and three studies.

## 4 Discussion

Our AdjMaxP method is a simple approach for identifying conserved statistical associations of shared features across omics studies under inter-study dependencies when unknown technical factors may limit the specificity of any one platform. Importantly, although for convenience, we distinguish between AdjMaxP and meta-analysis methods. AdjMaxP is similar to meta-analysis methods in that it is more sensitive than independently considering shared statistical significance across studies to infer conserved signals. Like Province-Borecki, AdjMaxP doesn’t require that all subjects be shared across studies or that the exact number of overlapping subjects be known (Province and Borecki, 2013).

Our simulation studies demonstrated that AdjMaxP should yield high precision under varying potential for technical artifacts in one or more studies. AdjMaxP consistently outperformed Province-Borecki in identifying conserved features under high levels of unconserved signal across studies. The performance of AdjMaxP was relatively unaffected by changes in the level of unconserved signal, whereas the performance of Province-Borecki was negatively affected by these changes. This is unsurprising, given that Province-Borecki is based on the sum of z-scores across studies. Thus, these tests can reach statistical significance if the signal from only one study is high.

Finally, several additional considerations are relevant when applying AdjMaxP. Since AdjMaxP significance testing is a function of the maximum p-value of a given feature across studies, we note that disproportionately higher weight is given to studies with less statistical power to detect genuine associations, either by greater noise and/or smaller sample size. However, this shortcoming is of no detriment to the overall precision but may negatively impact sensitivity for identifying these features. Additionally, spurious findings are possible if studies share features with technical artifacts, either by chance or by similar aspects of their molecular capture, e.g., multiple studies performed using similar sequencing or probe-based technologies. Thus, spurious findings will be limited when studies use disparate technologies.

In conclusion, AdjMaxP offers a novel strategy for improving the performance of biomarker discovery in non-independent cross-platform study designs. Moreover, given the potential for technical artifacts in the application of emerging high-throughput omics technologies, AdjMaxP promotes the adoption of such study designs for robust biological inference.

## Acknowledgements

This work was supported by the following National Institute on Aging (NIA) grants: UH2AG064704 (ER, PS), R01AG061844 (PS), U19AG023122 (PS).

