## Supplementary Figures for "A Simple Strategy for Identifying Conserved Features across Non-independent Omics Studies"

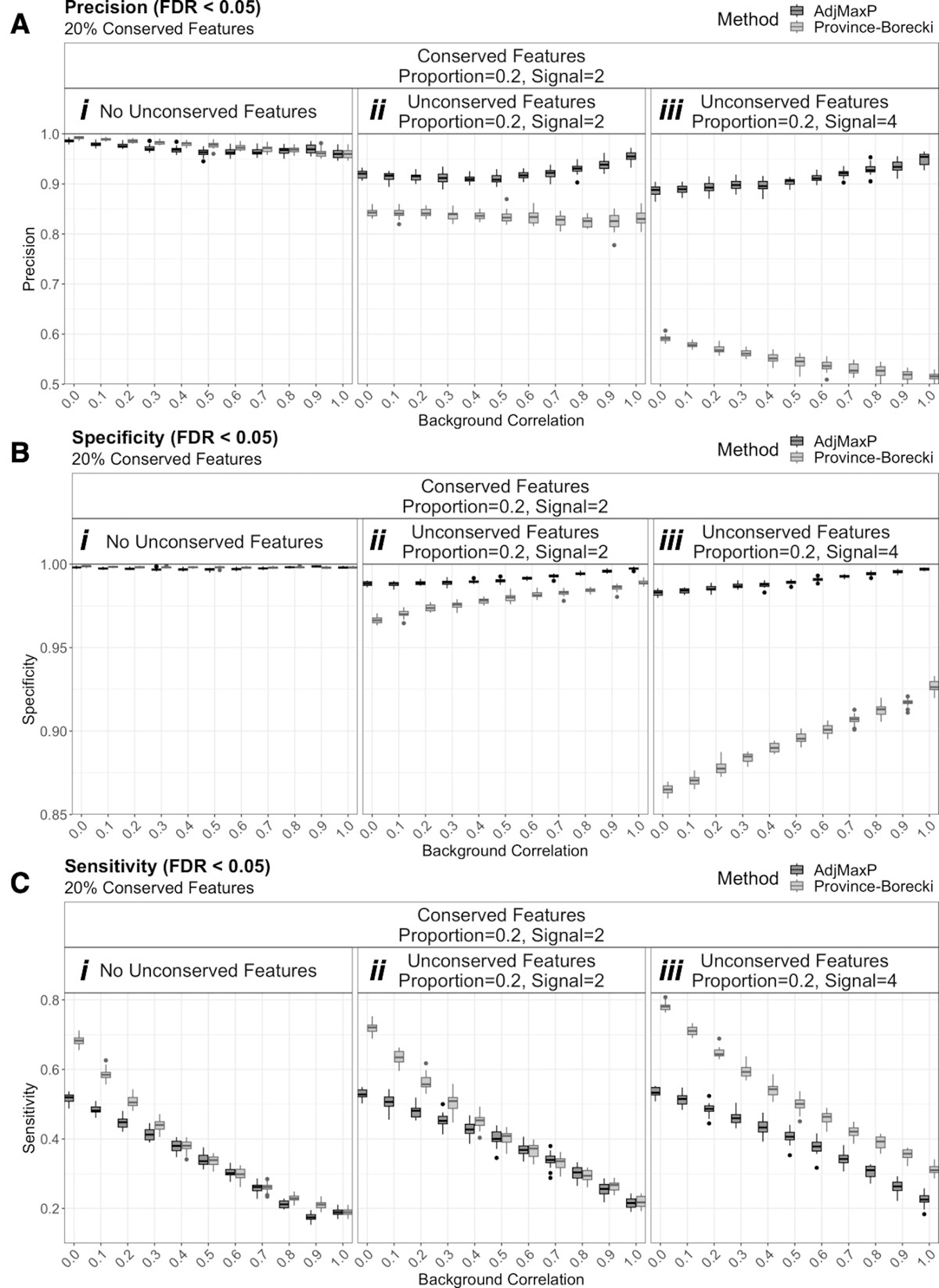

**Figure S1: Comparison of precision, specificity, and sensitivity from simulations of three studies with 20% conserved features, based on an FDR corrected q-value threshold of 0.05.**

Boxplots reflect the distribution of performance at FDR q-value threshold, 0.05, for 25 simulations of 10,000 shared features at different levels of background correlation and unconserved signal, across simulations of three studies with 20% conserved features. “Conserved” and “Unconserved” features refer to features for which signal was added (i.e. deviate from the null distribution) across all studies or fewer than all studies, respectively. “Proportion” and “Signal” indicate the proportion of features for which signal was added and the magnitude of added signal, respectively. Full simulation results are shown in Supplementary Table S2.

A) Precision
B) Specificity
C) Sensitivity

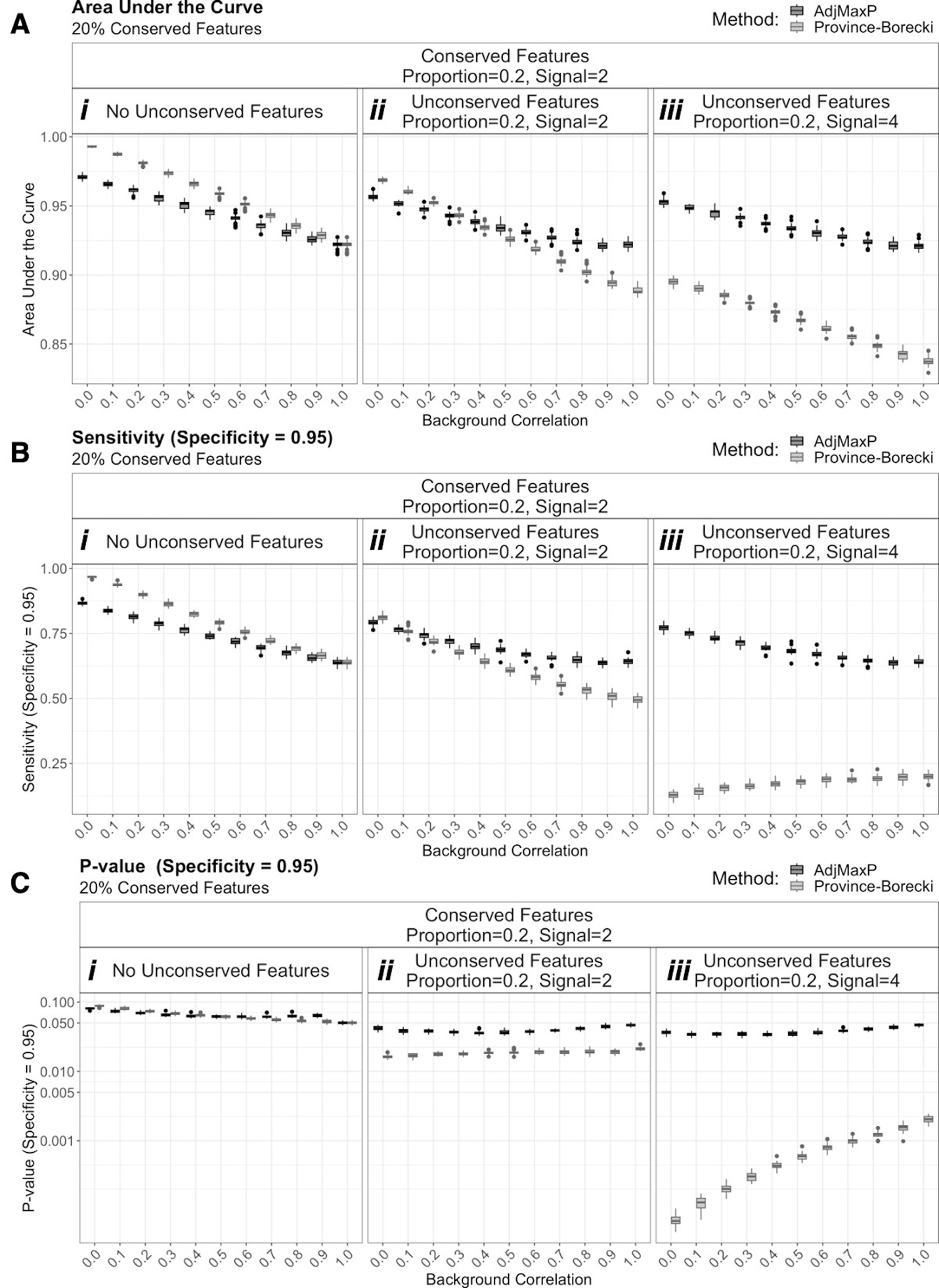

**Figure S2: Comparison of performance under threshold variable performance from simulations of three studies with 20% conserved features, based on an FDR corrected q-value threshold of 0.05.**

Boxplots reflect the distribution of performance for 25 simulations of 10,000 shared features at different levels of background correlation and unconserved signal, across simulations of three studies with 20% conserved features. “Conserved” and “Unconserved” features refer to features for which signal was added (i.e. deviate from the null distribution) across all studies or fewer than all studies, respectively. “Proportion” and “Signal” indicate the proportion of features for which signal was added and the magnitude of added signal, respectively. Full simulation results are shown in Supplementary Table S2.

A) Area under the curve.
B) Sensitivity of conserved feature identification when specificity is equal 0.95.
C) Nominal p-value threshold when specificity is equal to 0.95.
